# How nutrition and energy needs affect bumble bee pollination services: A mathematical model

**DOI:** 10.1101/2022.02.10.480001

**Authors:** Pau Capera-Aragones, Eric Foxall, Rebecca C. Tyson

## Abstract

The balance between nutrition and energy needs has an important impact on the spatial distribution of foraging animals. In the present paper, we focus on the case of bumble bees moving around a landscape in search of pollen (to meet nutritional needs) and nectar (to meet energy needs). Depending on the colony demands, bumble bees can concentrate their foraging effort towards either pollen rich flower species or nectar rich flower species. This behaviour allows us to establish a strategy – a spatial landscape design – which can maximize the pollination services of crops that are nutritionally deficient for pollinators by adding nutritionally rich wildflower patches. To do this, we formulate a mathematical partial integro-differential equation model to predict the spatial distribution of foraging bumble bees. We use our model to predict the location, composition and quantity of the wildflower patches adjacent to crop fields that will be most beneficial for crop pollination services. Our results show that relatively small quantities of wildflowers in specific locations with respect to the nest sites and the crop can have a positive impact on pollination services when the composition (i.e., pollen to nectar ratio) of the added wildflowers is significantly different from the composition of the existing crop flowers.

## 1 Introduction

Bumble bees are the predominant or exclusive pollinators of a large number of wild plants (Goulson, 2003). In addition, and most importantly for the economy, bumble bees provide important pollination services to many crops around the world (Corbet et al., 1991; Goulson et al., 2008, 2003; Kremen et al., 2002). Understanding the foraging behaviour of bumble bees is critical for the management of bumble bee mediated crops such as cucumber, pumpkin, watermelon, blueberry, and cranberry (Delaplane et al., 2000; Kremen et al., 2002; Richards, 2001). In places where managed honey bees (*Apis mallifera*) - the most commonly used pollinator to meet crop pollination requirements (Carreck and Williams, 1998) - are declining, a good understanding of the foraging behaviour of bumble bees is especially important. In the United States, for example, the critical services of honeybees is compromised by the decline in beekeeping, which has been estimated to be of around 50% during the last 50 years, and expected to decline further as the Africanized race of *A*.*mellifera* continues to spread (Kremen et al., 2002).

Buhk et al. (2018); Carvalheiro et al. (2011); Feltham et al. (2015) and Blaauw and Isaacs (2014) provide empirical evidence that the addition of wildflowers adjacent to a crop can lead to an increase in pollination services by wild pollinators such as bumble bees. However, a quantification of the location, quantity and type of wildflowers needed to optimize crop pollination is lacking, and so the need for a model is clear. Häussler et al. (2017) model crop pollination services and show that, due to an increase of pollinator population size, adding strips of wildflowers around crops can have a positive long-term effect (over multiple seasons) on crop pollination. However, in the same study, it is also said that adding strips of wildflower can have a negative short-term effect (within a single season) due to the competition for pollinators between flower species. Capera-Aragones et al. (2021) use a spatially explicit modelling approach to show that wildflower strips can be advantageous even in the short term if the appropriate location, quantity, and type of wildflowers are used. Their model includes more complexity in bumble bee foraging behaviour than previous work, particularly with respect to the influence of memory. In the present work, we are interested in understanding how the balance between nutritional and energy needs in bumble bee colonies changes the results predicted in Capera-Aragones et al. (2021).

During their foraging bouts, bees gather two different resources: nectar and pollen. Nectar provides water and sugar and is the main source of energy for bees, while pollen provides proteins, lipids, vitamins and minerals, and is important for rearing and development (Ball, 2007; Nicolson, 2011). Bumble bees therefore need to forage for both pollen and nectar. Different flower species have different compositions of nectar and pollen, hence offering different nutritional and energy resources. Thus, pollen and nectar foraging decisions will affect the rates at which the bees visit different types of flowers. These choices in turn, will affect the pollination services provided to the different flower species.

Free (1955) studied the division of labour within bumblebee colonies, including the division of foragers into nectar and pollen foragers. Free (1955) classified bees as pollen-foragers if they carry a pollen load in their corbiculae and as nectar-foragers if not. Pollen-foragers may also transport nectar, but nectar-foragers do not carry pollen. Bees showed constancy, i.e., bees that in the last bout collected pollen were more likely to collect pollen in the next bout. The proportion of pollen:nectar foragers is strongly related to the needs of the colony (Dornhaus and Chittka, 2005; Free, 1955; Kraus et al., 2019). Before a foraging bout, bees make an evaluation of the nectar and pollen stored in the nest, and then decide which resource to forage depending on which one is most limiting.

In this paper, we first expand the model in Capera-Aragones et al. (2021) to study how the division of labour between pollen and nectar foragers, and the effect of the interaction between the resource availability on the landscape and the resource needs of the colony, impact the dynamics of bee movement, foraging behaviour, and crop pollination services. We then show that our model can effectively reproduce the results of the empirical studies from Free (1955). At the end, we use our model to quantify the relationship between crop pollination services and the quantity, location, and relative pollen:nectar proportion of the wildflower patch added adjacent to the crop. It is important to note that in this work we do not include population dynamics, focusing instead on the effect of the spatial distribution and type of resources on the landscape. In particular, agricultural landscapes tend to have plentiful monotype resources from a crop for a few weeks, followed by many more weeks of much less abundant but mixed wildflower resources. In our work, we find that pollen-rich wildflower patches planted near pollen-deficient crops can result in increased pollination services to the crop, even in the absence of any demographic effect, if the wildflowers are planted in the correct location and provide the necessary complement to the crop resources.

## 2 Model and Methods

We begin with a brief overview of previous work upon which our model is built. In Tyson et al. (2011) a spatially-explicit model of honey bee dispersal is presented. In this model, the foraging population is divided into two sub-populations, one engaged in an intensive search mode (bees that are actively harvesting on resource) modeled using diffusion, and the other engaged in an extensive search mode (bees that are scouting for new resources to harvest) modeled by advection. In the same paper, it is shown that dividing the population into two sub-populations provides a better fit to organism movement data than models in which the population is considered homogeneous. Based on Tyson et al. (2011), Capera-Aragones et al. (2021) recently developed a mathematical model for wild bees, in which the foraging population is split into three different interacting sub-populations:

- Harvesting bees (*H*): those individuals that are collecting floral resources on the landscape and moving randomly from one flower to another (by diffusion).
- Scouting bees with memory (*S*): those bees that have knowledge of the landscape and use directed flight (advection) to move efficiently over larger distances, from one patch of flowers to another, or from the nest to a patch of flowers. These bees can be identified as exploiters if we follow the notation used in the empirical work done by Woodgate et al. (2016).
- Scouting bees without memory (*Sn*): those bees that do not have knowledge of the landscape and use directed flights (advection) as well as diffusion to explore the landscape and find new patches of flowers. These bees are identified as explorers in Woodgate et al. (2016).

as discussed below, the division into three sub-populations is necessary for a detailed description of the foraging distribution of of wild bees with memory. In the present paper, we also divide foraging bees into these three sub-populations performing different movement modes.

Similarly to Capera-Aragones et al. (2021), in our work, model bees move between the mentioned sub-populations at rates that depend on both resource density and the density of bees of each type. For example, scouting bees with memory at location *x* are likely to become harvesters at *x* as the floral resource density at *x* increases. The density of resource in the landscape affects the transitions between bee sub-populations, and the sub-populations affect the density of resource in the landscape via resource depletion.

Bees are required to return to the nest to unload the resource collected after each foraging bout. That is, they are central-place foragers. In contrast to previous work from Moorcroft et al. (2006) and Capera-Aragones et al. (2021) where the central-place foraging behaviour is included by adding a constant bias (advection) towards the nest to the foraging population, in the present work, we model the central-place foraging by assuming that harvesters are continually teleported to the nest at a constant rate (see Eq. (2.1)). Teleporting is used because it implies that harvesters reach the nest at a specific moment in time (which is not the case with a linear advection term), at which time bees can decide between becoming a pollen or a nectar scouting bee, depending on the pollen and nectar storage in the nest.

One of the key contributions in Capera-Aragones et al. (2021) is the inclusion of memory effects in a spatial differential equation model for bee foraging. Memory is an important tool used by bees to improve their foraging efficiency. Based on findings of radar tracking studies of bees (Lihoreau et al., 2012, 2013; Woodgate et al., 2016, 2017), Capera-Aragones et al. (2021) include memory by making the sub-population of scouting-bees with memory to be more likely to fly advectively in a chosen flight direction. That is, those bees with knowledge of the landscape are less likely to use direct motion (advection) on a random direction, and more likely to use the direct motion towards those locations where they remember finding resources. By contrast to other recent approaches such Shi et al. (2019a,b) and Oliveira and Berbert (2020), in this work we use the one developed in Capera-Aragones et al. (2021).

The main difference and contribution of this paper with respect to the model in Capera Aragones et al. (2021) is the division of food into two different variables: pollen and nectar, with the consequent division of the bee population into two: pollen foragers and nectar foragers. This division means that the new model doubles the number of sub-populations. Now, foraging bees can be in six different states instead of only three: pollen harvesters (*H*^*P*^), pollen memory scouts (*S*^*P*^), pollen no-memory scouts (*Sn*^*P*^), nectar harvesters (*H*^*N*^), nectar memory scouts (*S*^*N*^) and nectar no-memory scouts (*Sn*^*N*^). Fig. 1 is the compartmental diagram of the model presented in this paper.

**Figure 1:**
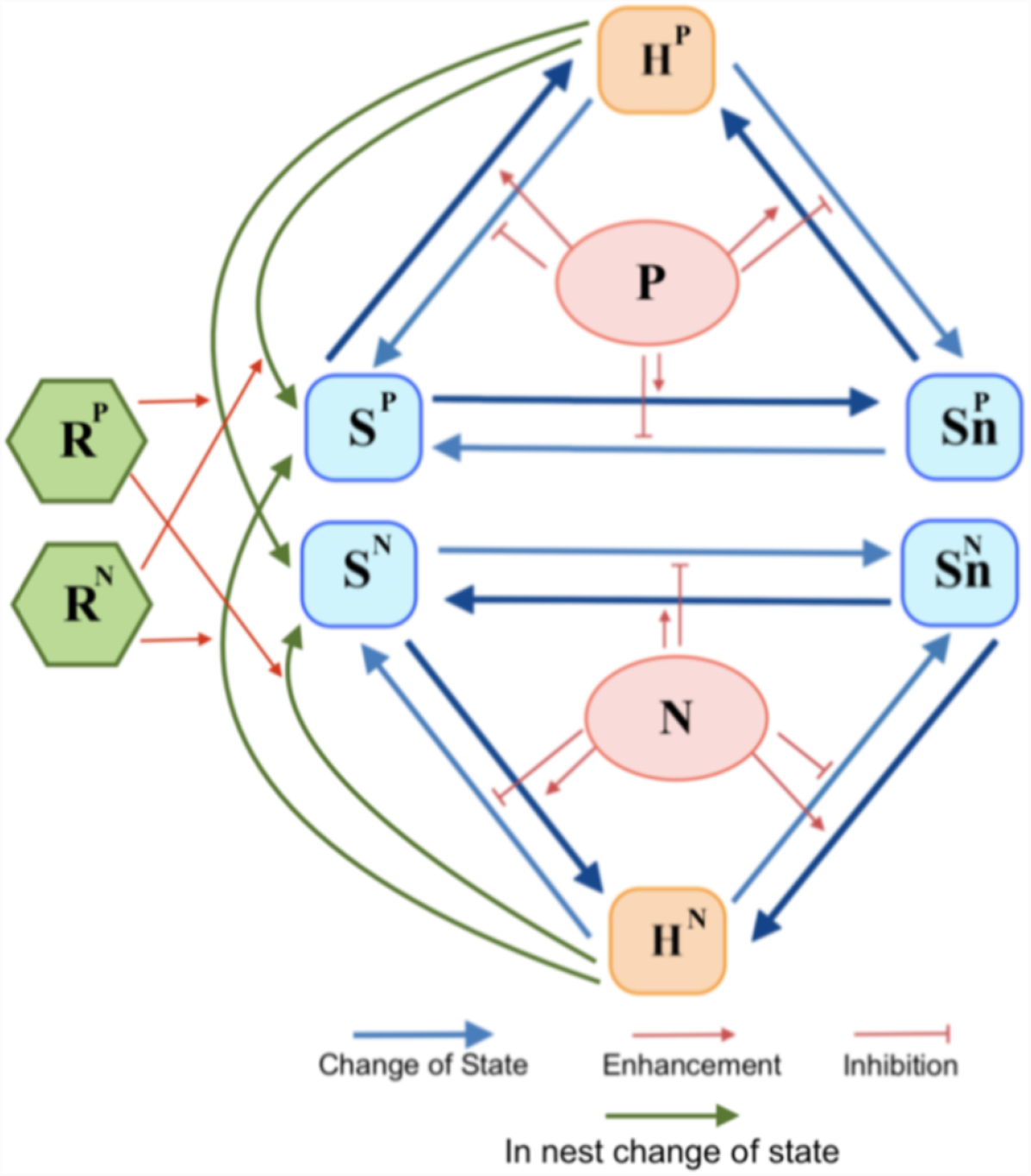
Compartmental diagram of the model under consideration. Our model splits bees into six sub-populations depending on their movement modes and their foraging objective: pollen harvesters (*H*^*P*^), pollen scouting bees with memory (*Sn*^*P*^), pollen scouting bees without memory (*Sn*^*P*^), nectar harvesters (*H*^*P*^), nectar scouting bees with memory (*Sn*^*P*^), nectar scouting bees without memory (*Sn*^*P*^) (each of *S*^*P/N*^ and *Sn*^*P/N*^ have left and right moving sub-populations). Pollen foragers (*H*^*P*^, *S*^*P*^ and *Sn*^*P*^) are able to change between movement modes and the transitions are enhanced or inhibited by the presence of pollen (*P*), equivalently for nectar foragers with the presence of nectar (*N*). The sub-populations of total pollen foragers and nectar foragers interact through the pollen and nectar resource stored in the nest (*R*^*P*^ and *R*^*N*^).

For simplicity, we assume that pollen foragers (including pollen harvesters, pollen memory scouts and pollen no-memory scouts) do not interact with nectar foragers (nectar harvesters, nectar memory scouts and nectar no-memory scouts) outside the nest. That is, once a forager has decided to harvest pollen, it cannot become a nectar forager (or vice-versa) until its current foraging bout has finished. This resource specialisation (to either pollen or nectar, broadly, not type of pollen or nectar) during a single foraging bout is based on the empirical results found in Free (1955) and Hagbery and Nieh (2012). Following this empirical evidence, the interaction between pollen and nectar foragers occurs only in the nest at the beginning of each foraging bout. The interaction is such that nectar foragers are likely to become pollen foragers if the proportion of pollen stores relative to the pollen requirements of the colony (*θ*^*P*^*R*^*P*^) is smaller than the proportion of nectar stores relative to the nectar requirements of the colony (*θ*^*N*^*R*^*N*^) and are likely to continue as a nectar forager if the colony’s needs are reversed. Similarly, former pollen foragers may or may not switch to nectar foraging at the beginning of a new foraging bout.

Our model considers the time evolution of the pollen stored (*R*^*P*^) and nectar stored (*R*^*N*^) in the nest to be a function of the pollen/nectar collected by harvesters and the pollen/nectar consumed by all bees.

Mathematically, we can express the model presented in this paper by the set of partial integro-differential equations (2.1):

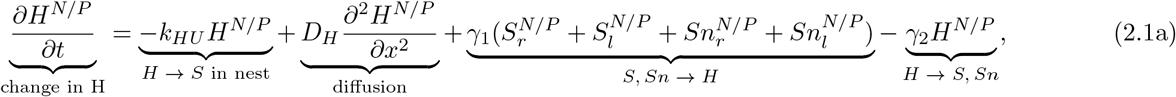

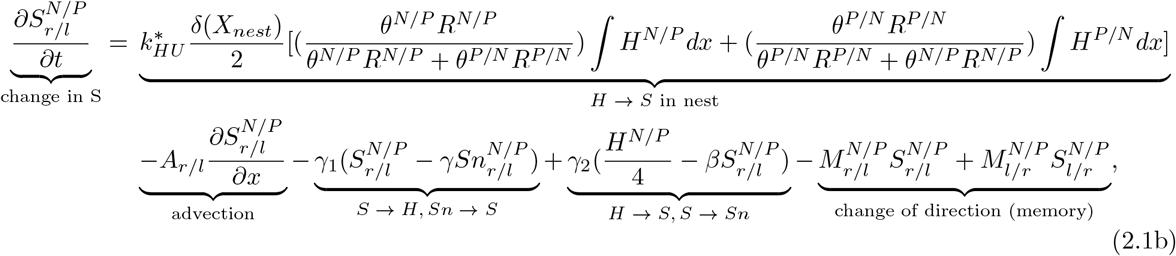

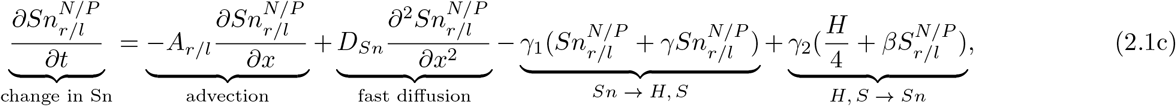

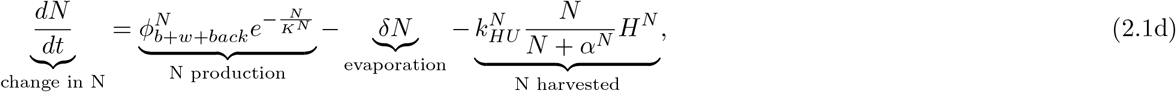

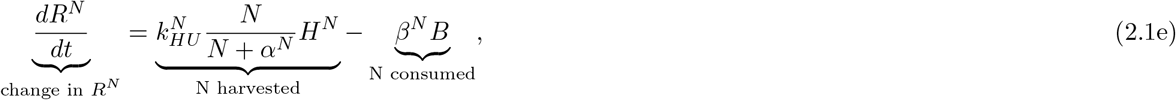

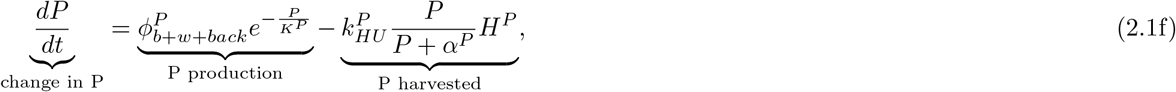

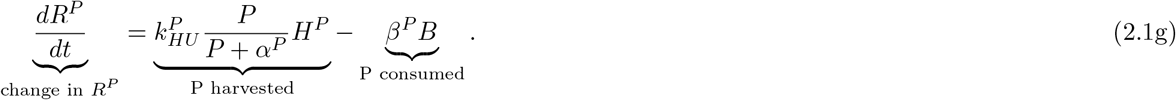

where all parameters are defined in the Table. 2. 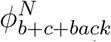 represents the nectar production at time *t* and position *x* by crop flowers (*b*), wildflowers (*w*) and background flowers (*back*), all togheter. 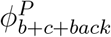 is equivalent to 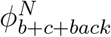 but for pollen production. 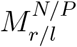 are the memory functions which are defined for each pollen and nectar forager independently and take the same form as in Capera-Aragones et al. (2021):

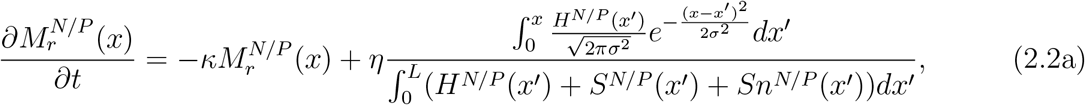

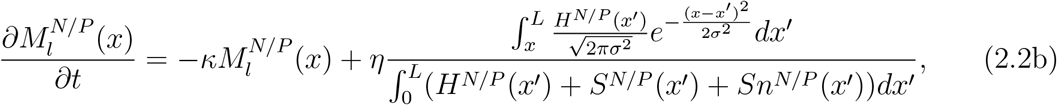

where *κ* represents the decay rate of memory over time and *σ* the standard deviation of the spatial memory kernel. 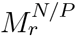 and 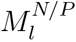 are, respectively, the rates at which a left/right-moving scout switches direction (i.e., becomes right/left-moving).

**Table 1:**
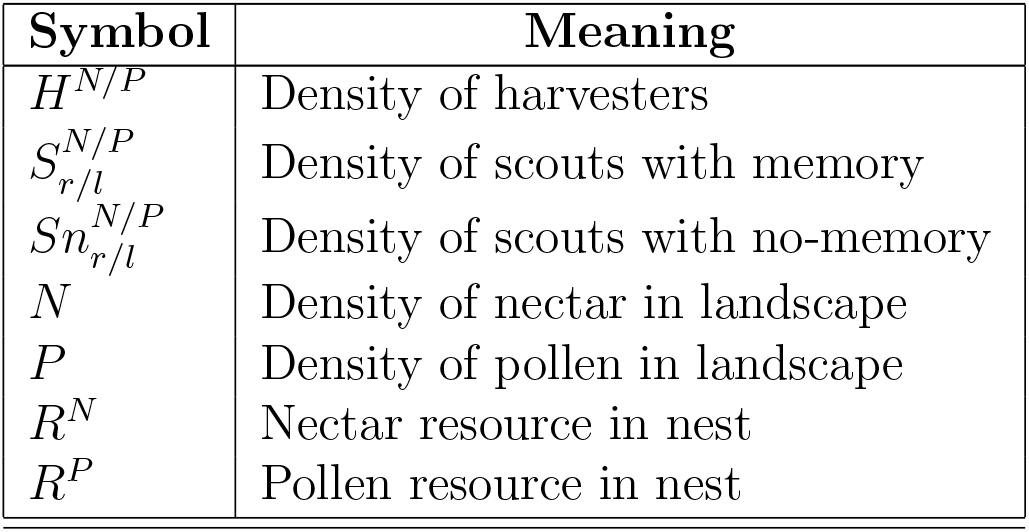
Description of state variables of the model. The superscripts *N* and *P* refer to nectar and pollen foragers respectively, and the subscripts *r/l* on *S* and *Sn* refer to right and left moving scouting bees.

| Symbol | Meaning |
| --- | --- |
| $H^{N/P}$ | Density of harvesters |
| $S_{r/l}^{N/P}$ | Density of scouts with memory |
| $Sn_{r/l}^{N/P}$ | Density of scouts with no-memory |
| $N$ | Density of nectar in landscape |
| $P$ | Density of pollen in landscape |
| $R^N$ | Nectar resource in nest |
| $R^P$ | Pollen resource in nest |

**Table 2:**
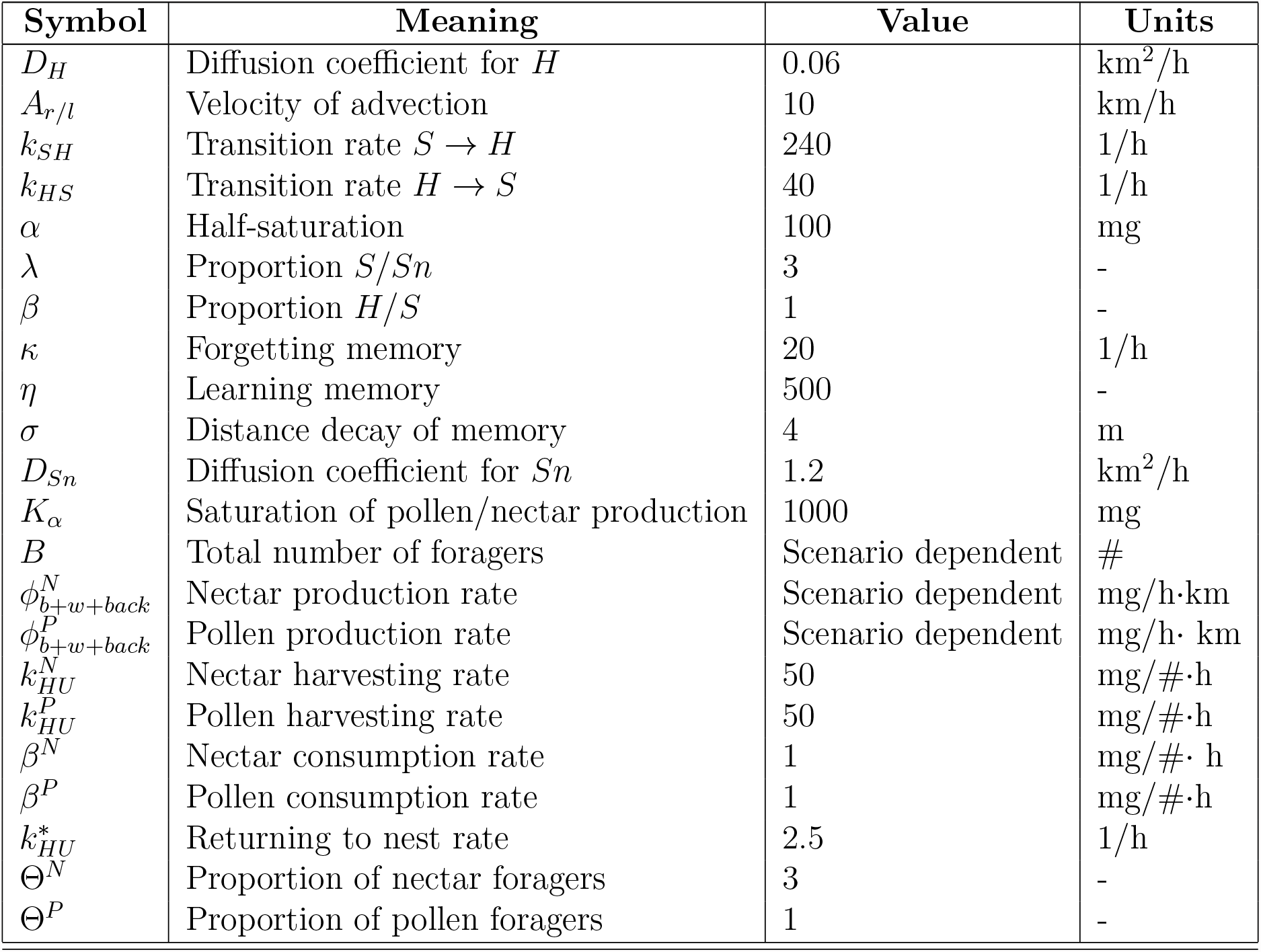
Description of the parameters of the model. The superscripts *N* and *P* refer to nectar and pollen foragers respectively. All parameters values are derived in our previous work (see (Capera-Aragones et al., 2021)), except for the last five. 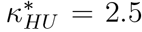 is the value that limits foraging range at 2km from the nest site. Θ^*N*^ = 3, Θ^*P*^ = 1, *β*^*N*^ = 0.001 and *β*^*P*^ = 0.001 are the values that fix the proportion of *R*^*N*^ and *R*^*P*^ according to the data in Free (1955).

The nectar/pollen enhancement/inhibition switching rates take the same form as in Capera-Aragones et al. (2021),

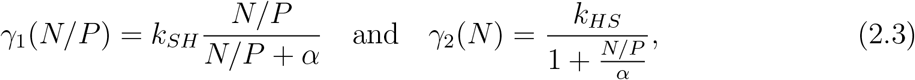

where *k*_*SH*_ give the maximum transition rate from scouting to harvesting in the presence of *N* or *P*, and *α* is the half-saturation constant that determines when the switching rates *γ*_1_ and *γ*_2_ are strongly or weakly resource dependent. The parameter *k*_*HS*_ gives the maximum transition rate from harvesting to scouting.

### 2.1 Metrics

In this section we define the metrics that we will use to describe and quantify the output of our model. Apart from measuring the number of harvesting bees, scouting bees with memory and scouting bees without memory on pollen and nectar (*H*^*N*^, *H*^*P*^, *S*^*N*^, *S*^*P*^, *Sn*^*N*^, *Sn*^*P*^), we are interested in measuring the resource collected (pollen and nectar) across the landscape, which we consider to be proportional to the pollination services. We track the resource collected (pollen and nectar) from two different types of flower species (crop flowers and wildflowers) as a function of location and time (*x, t*):

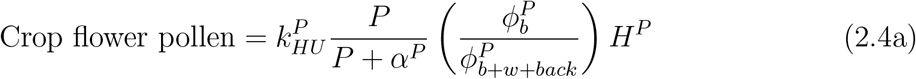

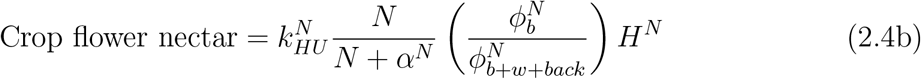

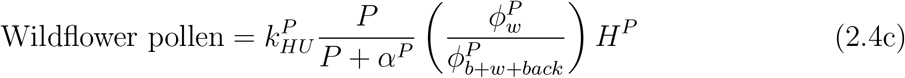

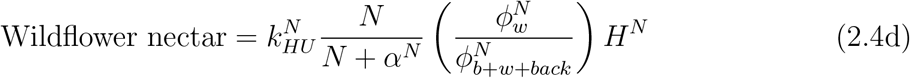

where 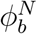 and 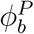 are the nectar and pollen produced by crop flowers respectively, and 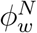and 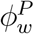 are the nectar and pollen produced by wildflowers, and 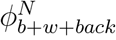 and 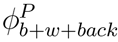 are the nectar and pollen produced by crop, wildflowers and background flowers all together. Using eqs. (2.4), we can determine the quantity of pollen and nectar collected from crop flowers and wildflowers in a specific location at a specific time. We can also compute it over the entire landscape (integrating with respect to space) at a particular time, or during a specific time interval (integrating with respect to time). Note that in eq. (2.4a), the term 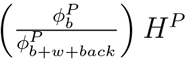 refers to the amount of pollen harvesters which are collecting pollen from crop flowers, and 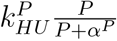 refers to the saturating dependence of resource collection, similarly for the other cases in eqs. (2.4).

### 2.2 Numerical approach

The model described consists of eighteen partial integro-differential equations (one for each of the state variables 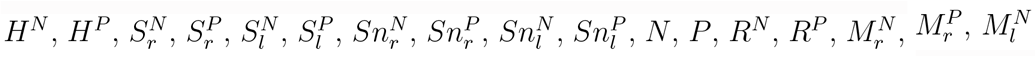 and 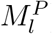), coupled with each other. The initial conditions consist of

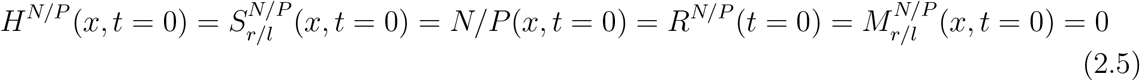

and

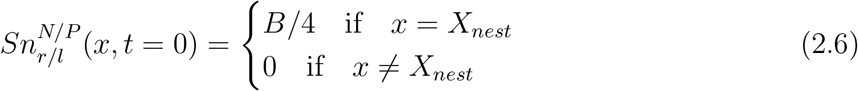

We applied no-flux boundary conditions to all the state variables involving movement:

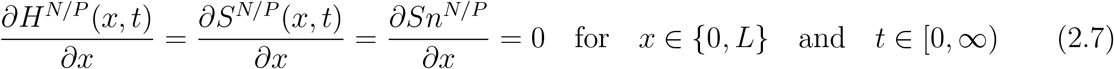

To solve the system of equations we used a fractional step method to decouple the distinct modelling processes in Eqs.(2.1), and the best suited numerical method for each of the processes was applied sequentially (see Tyson et al. (2000)). In a first stage we solved the diffusion and advection terms using an implicit TR-BDF2 method (Hosea and Shampine, 1996) and in a second stage we solved the non-linear reaction terms using an explicit RK4 method (La et al., 2002). To solve the integrals in Eqs.(2.2) we used the Simpson Rule. The spatial step used was 50m and the time step 1 second.

## 3 Results

### 3.1 Effect of the division of food into pollen and nectar on foraging spatial distributions

In this section we show how the new mechanism added in model Eqs. (2.1) to account for the balance between nutrition and energy needs of foragers produce the characteristic behaviours of bumble bees observed in Free (1955).

First, we investigate the changes in the proportion of nectar and pollen foragers as a consequence of changes in landscape resource availability. Fig. 2 (left), we show that the higher the proportion of pollen:nectar resource availability in the landscape, the smaller the proportion of pollen:nectar foragers in the colony. This inverse proportion occurs because fewer bees are needed to gather the necessary quantity of the easily accessed resource.

**Figure 2:**
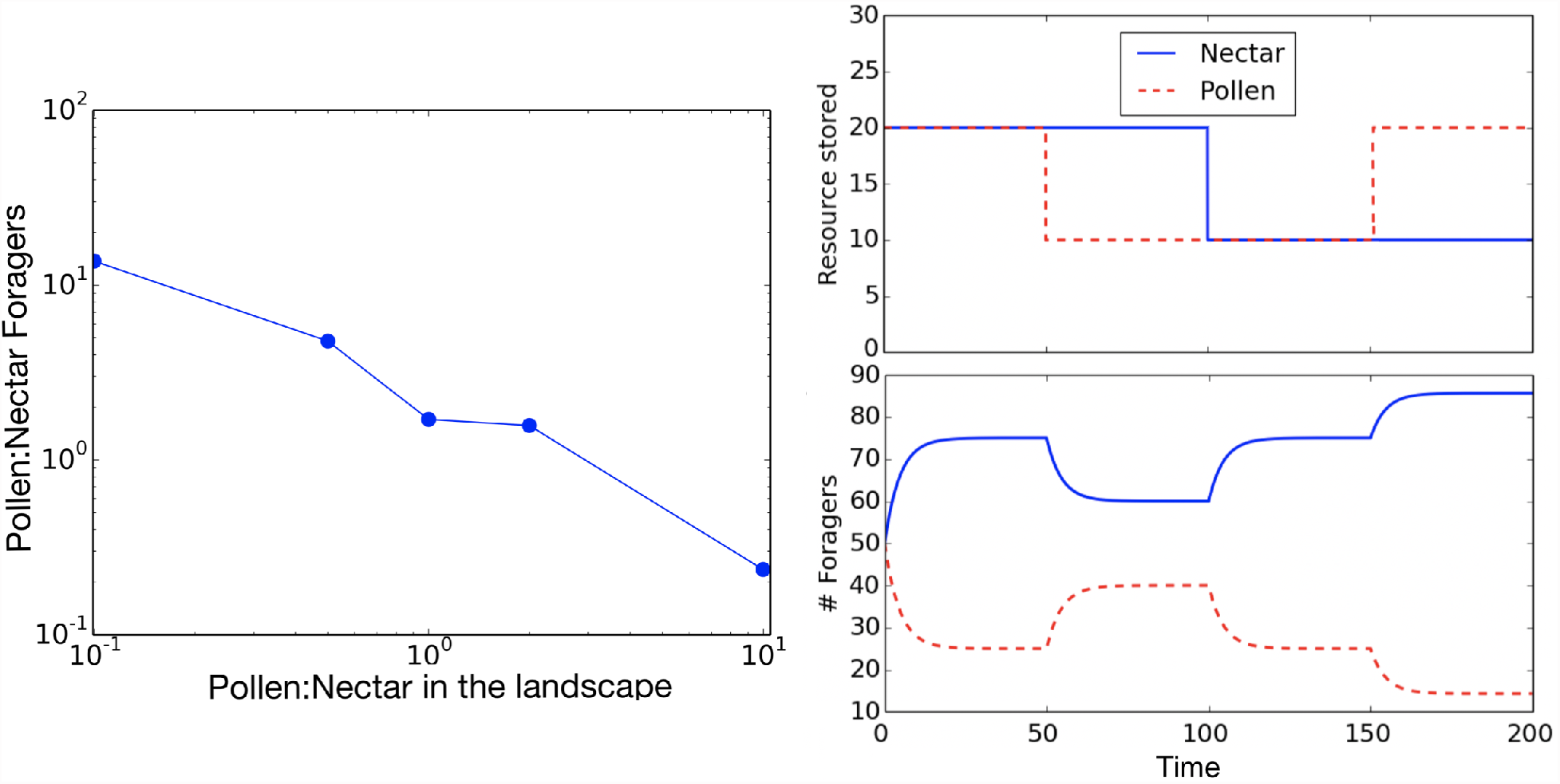
Pollen and nectar foragers as functions of resources on the landscape (left) and resources in the nest (right). On the right, nest resources are fixed at the values shown (top), and the foraging population is allowed to respond (bottom). The four storage scenarios investigated on the right are: (1) equal (high) amounts of nectar and pollen (*t* = (0*h*, 50*h*)), (2) twice as much nectar as pollen (*t* = (50*h*, 100*h*)), (3) equal (low) amounts of nectar and pollen (*t* = (100*h*, 150*h*)), and (4) twice as much pollen as nectar (*t* = (150*h*, 200*h*)).

Second, we examine the foraging behaviour responses to “in nest” nectar/pollen storage depletion. In Fig. 2 (right), we use our model to show how the numbers of pollen and nectar foragers depend on the quantities nectar and pollen stored in the nest. To obtain the results in Fig. 2 (down right) we held the stored resource values *R*^*P*^, *R*^*N*^ constant at the values indicated in Fig. 2 (top right). For the rest of this paper, we do not fix the resource stored in nest, instead it increases or decreases over time according to Eq. (2.1).

### 3.2 Wildflower plantings benefit crop pollination

In this section, we evaluate how the balance between nutritional and energy needs affects the spatial distribution of bees. In the presence of flower species with different pollen/nectar proportions, bees can be strongly biased towards the location of one flower species or another depending on the colony’s needs.

The landscapes under consideration in this section will always consist of a crop and a nesting site at fixed locations, a wildflower patch and a uniform low-density wildflower matrix (wildflower background). See Fig.3 (top) for a depiction of the linear landscape under consideration. In order to study the possible benefits of a wildflower patch enhancement on crop pollination services, we will look at the pollination services as a function of four different variables:

**Figure 3:**
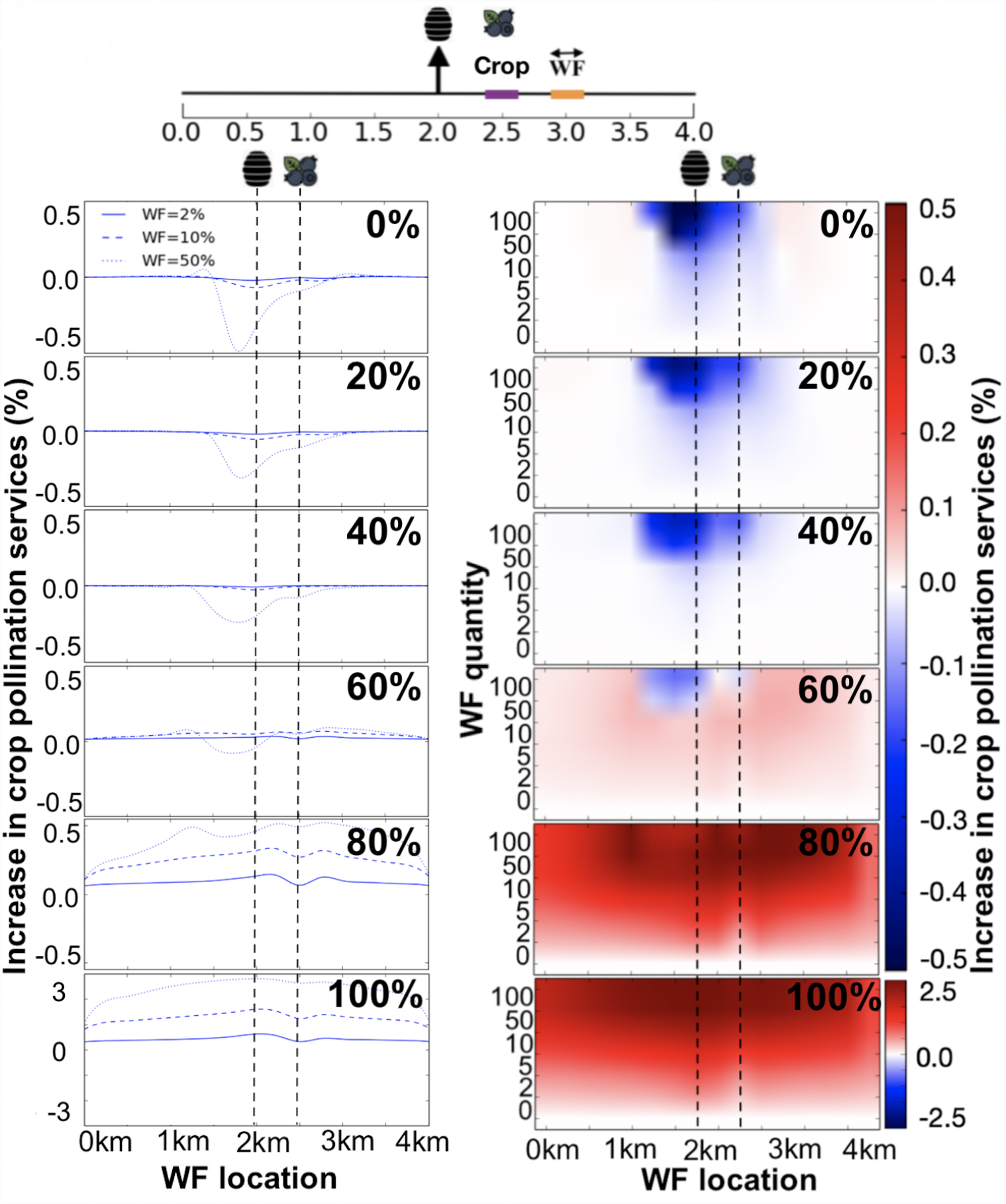
Top: Layout of the linear landscape. It consists of a nest in the middle (vertical arrow), a crop patch of width 0.5 km (thick purple segment, colour online) and a wildflower patch enhancement (thick orange segment), which can be placed anywhere in the landscape (double headed arrow). There is also a low density wildflower background across the entire landscape. Left six panels: Crop pollination services as a function of the location of the wild flower patch enhancement for three different wildflower patch densities (WF = 2%, 10% and 50%), and for six different wildflower differentiation values (Eq. 3.2). Right six panels: Crop pollination services as a function of the location and quantity of the wild flower patch enhancement, for six different wildflower differentiation values (Eq. 3.2). The locations of the bumblebee nest and crop are fixed and indicated by the dashed lines. Red (blue) colours indicate an increase (decrease) of pollination services in the crop. Note that the vertical scale is not linear, and the bottom plot has a different colour scale.

- Wildflower patch location: The position of the wildflower patch is defined in a 4 km landscape where there is a nest site fixed at position *l* = 2 km and a crop of width 0.5 km centred at *l* = 2.5 km.
- Wildflower patch quantity: The quantity of resource in the wildflower patch is defined relative to the quantity of resource in the crop. For example, a wildflower patch quantity of 10% means the total amount of resource in the wildflower patch is 10 times smaller than that in the crop.
- Wildflower patch differentiation (type): We define “wildflower differentiation” in relation to the crop. That is, we wish to define a metric where a wildflower differentiation of 100% means that the wildflower resources are completely different in pollen and nectar composition from the crop resources, while a wildflower differentiation of 0% means that the compositions are exactly the same. We thus define wildflower differentiation as a percentage given by

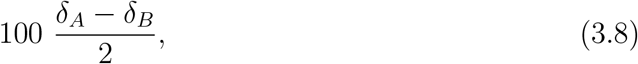

where

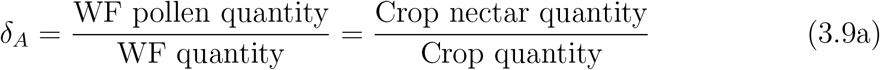

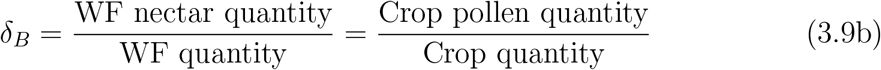

and *δ*_*A*_ and *δ*_*B*_ range between 0 and 2 and satisfy *δ*_*A*_ + *δ*_*B*_ = 2. We are assuming that the wildflower pollen quantity relative to the wildflower quantity is the same as the crop nectar quantity relative to the crop quantity, and vice versa (where “nectar” and “pollen” are swapped). This assumption enables a simpler interpretation of the results. Using Eqs. (3.8) and (3.9), wildflower differentiation of 100% corresponds to (*δ*_*A*_, *δ*_*B*_) = (2, 0) or (0, 2). In the first case, wildflowers only produce pollen whereas crop flowers only produce nectar. The second case represents the opposite scenario. In contrast, wildflower differentiation 0% occurs when (*δ*_*A*_, *δ*_*B*_) = (1, 1) and both wildflowers and crop flowers produce the same proportion of pollen and nectar.
- Wildflower background density: In addition to the crop field and the added wildflower patch, the landscape under consideration also includes a low density uniform distributed wildflower background. The density of the wildflower background is a measure of agricultural intensity where low densities represent intensively farmed landscapes with few wildflower resources other than the crop. We can measure the wildflower background density by comparing its integral over all the landscape to the quantity of resource in the crop.

Fig.3 shows the crop pollination services as a function of the wildflower patch location (left plot) and the crop pollination services as a function of the wildflower patch location and wildflower patch quantity (right plot). We show the pollination services for six different wildflower differentiation values, ranging from 0% to 100%.

Regarding the location of the wildflower patch enhancement, our results in Fig.3 show that planting the wildflowers in a location such that the crop is between the wildflowers and the nest site (wildflowers at *l ≈* 3km, where the nest is at *l* = 2 km and the crop is at *l* = 2.5 km) is a consistently good location regardless of changes in the wildflower differentiation and wildflower quantity. We can also observe that wildflowers planted close to the nest site (*l ≈* 2km) can lead to the highest increase in crop pollination services if the differentiation is large (“differentiation” *≈* 100%), but lead to the highest decrease in crop pollination services if the differentiation is low (“differentiation “*<*100%). Planting the wildflowers in the same location as the crop (*l* = 2.5km) appears to be disadvantageous in the majority of cases, whereas planting the wildflowers between the crop and the nest site appears to be in all cases less advantageous than when the wildflowers are planted such that the crop is between them and the nest site.

Regarding the quantity of the wildflower patch enhancement, we observe that when the differentiation is large (“differentiation” *>*80%) higher quantities are always more advantageous than lower quantities. By contrast, for smaller differentiation values (“differentiation “*<*80%) intermediate quantity values are the most advantageous, with high quantities located close to the nest site being the most disadvantageous and small quantity values located in the far edge of the crop with respect to the nest being the most advantageous.

With respect to the differentiation of the wildflower patch enhancement, the tendency shown in Fig.3 is clear; the higher the differentiation, the higher the increase in pollination services associated with planting wildflowers.

Fig.4 shows how the wildflower background affects the predictions in Fig.3. In Fig.4 we can see that, the smaller the wildflower background density (the more intense the agriculture), the larger the benefits of adding a wildflower patch to crop pollination services.

**Figure 4:**
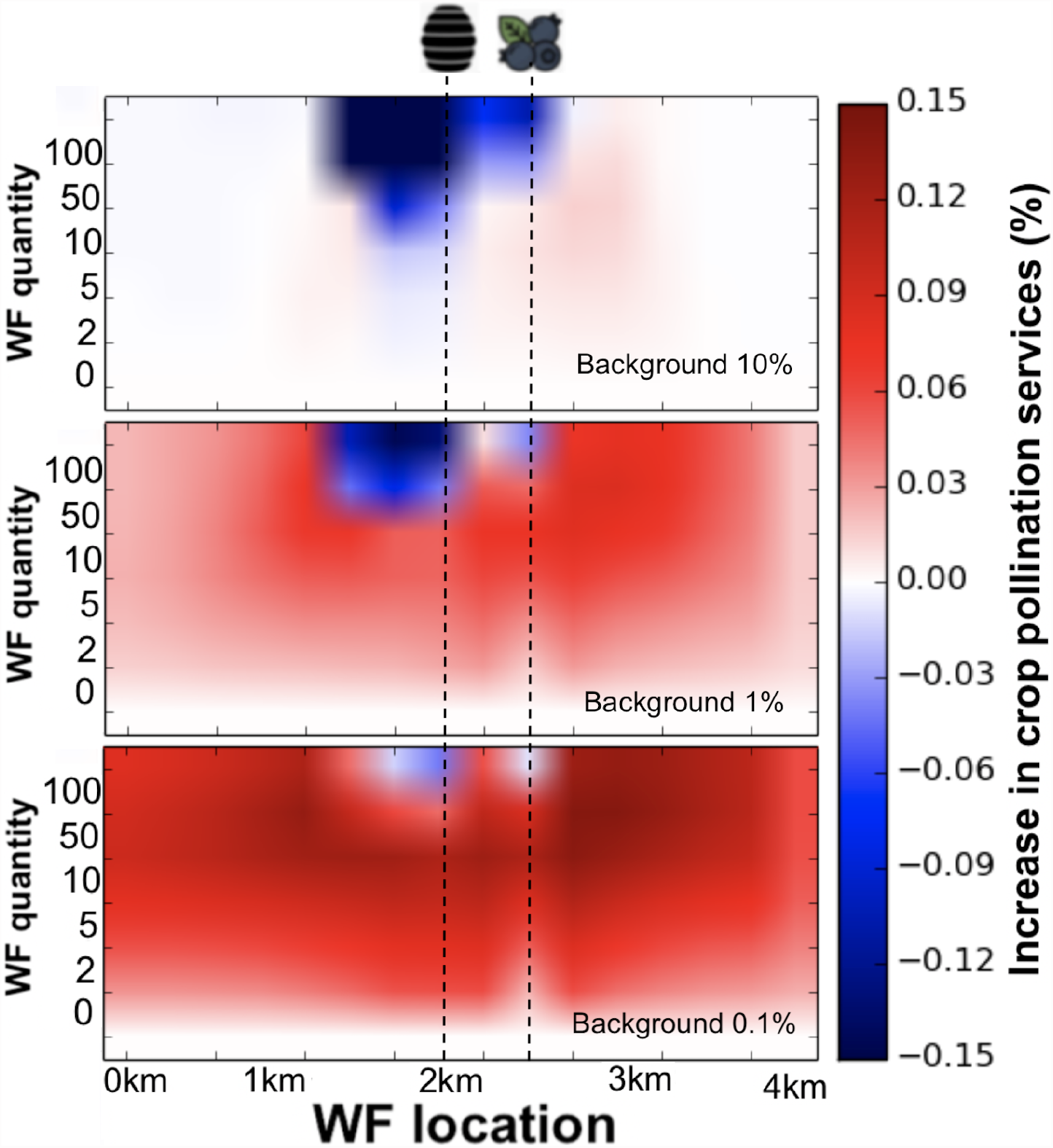
Crop pollination services as a function of the location and quantity of the wildflower patch enhancement for the case of wildflower differentiation (type) 90%. The locations of the bumble bee nest and crop are indicated by the dashed lines, respectively. The remainder of the landscape is covered by a low density wild flower background (see Fig. 3). The three panels show the effect of the density of the wildflower background on crop pollination services. Darker colour indicates higher pollination services in the crop. Note that the colourbar scale varies from one subplot to the next. As with Fig.3, crop pollination services are assumed to be proportional to the sum of the pollen and nectar collected.

Fig.5 shows the increase in crop pollination services as a function of the wildflower background density for three different wildflower patch densities and three different wildflower patch differentiation. Our results highlight that:

**Figure 5:**
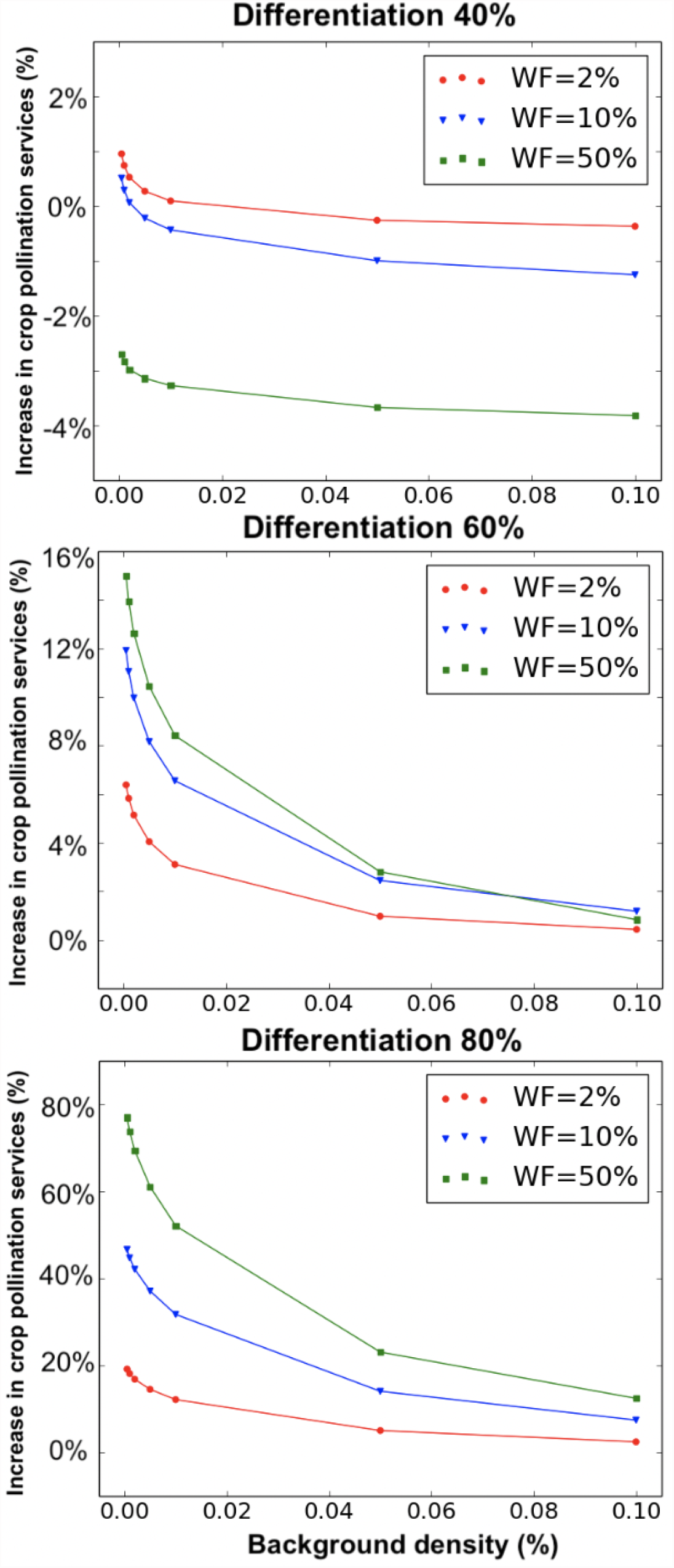
Increase in crop pollination services as a function of the wildflower background density for three different wildflower types (differentiation) and quantities. In this figure, the location of the wildflower patch is always in the far edge of the crop, at *l* = 3km.

- The greater the wildflower differentiation value, the larger the benefits of adding a wildflower patch. The increase in pollination services can vary between 0% to more than 80%.
- The more dense the wildflower patch, the higher the benefit, though for wildflowers very similar to the crop (low differentiation), smaller wildflower densities give better results (see Fig. 5 top).
- The smaller the density of the wildflower background, the larger the benefit of planting wildflowers. That is, the more intense the agriculture, the greater the benefit of planting a wildflower patch.

For the simulations in Fig. 3, Fig. 4 and Fig. 5, the total number of foragers is *B* = 6000 and the total amount of resource produced by the crop is 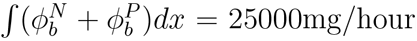. The total amount of resource produced by the wildflower patch and wildflower background is scenario dependent. It is important to note that rather than the absolute value of *B* or 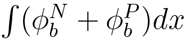, what changes the output of our simulations is their relative values, i.e., how big is the crop with respect to the existent population of bees in the landscape. In this work, the relative values are chosen such that a significant fraction of the resources are consumed.

## 4 Discussion

In this paper we expand the model presented in Capera-Aragones et al. (2021) to incorporate the effects of the balance between the two floral resources, i.e., pollen (nutrition) and nectar (energy), on bumble bee spatial distributions. We divide the forager population into two groups, pollen foragers and nectar foragers, and assume that foragers switch between groups depending on the colony needs. This assumption is consistent with Free (1955) and with the observation that colonies with a large amount of brood forage more often for pollen than colonies with a smaller brood population because of increasing pollen needs associated with brood development (Kraus et al., 2019).

In addition to the modelling of the two resource types and the two foraging populations associated to them, in the present paper, we applied central-place foraging behaviour using a different method than in Capera-Aragones et al. (2021). It is relevant to note that the different method does not change the model predictions significantly if correct parameter values are used. The method used in the present paper assumes that harvesters stop foraging (and return to the nest) at a constant per capita rate. The result is a decrease in the harvesting population. In order to keep the population of bees constant, we add a source of scouting bees at the nest location so that the number of scouts emerging at the nest location is equal to the number of harvesters disappearing from the landscape. In other words, we assume that harvesting bees become scouts in the nest, and ignore the travel time. The advantage of using this approach to model central place foraging behaviour is that it enables foragers to constantly decide between becoming a pollen or a nectar scouting bee, depending on the pollen and nectar storage in the nest (see Eq. (2.1)), thus allowing us to easily incorporate the effects of the balance between nutrition and energy needs.

### 4.1 Effects of the division of food into pollen and nectar

Free (1955) and Vaudo et al. (2015), find that bumblebees forage for pollen or nectar in proportion to the needs of the colony. In our work, we consider a relatively short period of time during which we can assume a constant population and constant colony needs. Based on Free (1955) and Vaudo et al. (2015), we assumed that bumble bees forage for pollen and nectar with the goal of storing three times as much nectar as pollen. As a consequence, we found the proportion of pollen/nectar foragers to be inversely proportional to the proportion of pollen/nectar resource in the landscape, but insensitive to the total amount of resource available (see Fig.2). Further work is needed to determine whether it is simply the proportion of each type of resource that matters, or if it is also the total quantity of resource, or some combination of these two measures.

In Fig. 2 (right), we use our model to reproduce some of the experiments in Free (1955), in which the reserves of pollen or nectar in nest are depleted and a subsequent increase of pollen or nectar foragers are respectively observed. These results provide qualitative support for our model.

### 4.2 Wildflower plantings benefit crop pollination

In our earlier work (Capera-Aragones et al., 2021), we showed that adding intermediate wildflower quantities in a location such that the crop is between the nest site and the wildflowers lead to the largest increase in crop pollination services. In the present paper, in addition to the location and quantity of wildflowers, we investigated the effects of the pollen and nectar composition of the wildflower patch (differentiation or type) relative to the crop and the effects of the density of the wildflower background (matrix) in the landscape.

Our results in Fig. 3 show that the greater the difference between the pollen/nectar composition of the wild and crop flowers, the larger the benefit provided by planted wildflowers to crop pollination services. In the differentiation 0% case (in which the composition of the wild and crop flowers is the same), the addition of wildflowers provides only a small benefit, and only if planted in small quantities and in the right location. If planted in large quantities and in a wrong location, the presence of wildflowers can significantly decrease the pollination services of the crop. This is due to competition between flower species for pollinators. In the differentiation 100% case (moment in which the resource composition of wild and crop flowers is completely different) the benefits are significant, independent of the location and quantity of wildflowers planted. It is relevant to note that, for the case in which the two flower species are completely different in composition, they never compete for pollinators and so, adding wildflowers never decreases the pollination services of the crop. In the case of wildflowers with an intermediate differentiation value (say, 10% to 90%), the interpretation of the results is harder to generalize since, depending on the location and quantities, adding wildflowers can lead to large increases or decreases in crop pollination services.

Fig. 4 and Fig. 5 show how the decrease of wildflower density in the background, which represents an increase in agricultural intensity, enhances the tendencies observed in Fig. 3. We can therefore conclude that intensively farmed landscapes stand most to benefit from the practice of planting wildflower patches adjacent to crops.

Looking at Fig. 3, Fig. 4, and Fig. 5 we arrive at the following conclusions regarding the benefits to crop pollination services of adding a wildflower patch:

- The more different in pollen/nectar composition is the wildflower patch with respect to the crop (high differentiation), the larger the benefits.
- Locating the wildflowers at the far side of the crop with respect to the nest site is consistently a good location with respect to changes in the “differentiation” or quantity of the wildflower patch enhancement, or changes in the density of the wildflower background. Locating the wildflower patch close to the nest site can be most beneficial if the “differentiation” is high (*≈*100%), but can be most detrimental for crop pollination if the “differentiation” is low.
- Large quantities of wildflower plantations are most beneficial if they have a high “differentiation” value, but small quantities are better if the “differentiation” value is small (see Fig. 5).
- The lower the density of the wildflower background, the larger the benefit of planting wildflowers. That is, the more intense the agriculture, the greater the benefit of planting a wildflower patch.

## 5 Conclusions

In this paper we investigated how the balance between nutrition (pollen) and energy (nectar) needs affect the spatial distribution of foraging bumblebees and the resulting pollination services they provide to a crop. We focused on a landscape consisting of two flower species (the crop and a wildflower patch) that can vary in pollen and nectar concentrations.

Our findings show that, in landscapes where pollen and nectar resources are not uniformly distributed, the balance between nutrition and energy needs can significantly affect the spatial distribution of foraging bumblebees. Our result is consistent with field studies indicating that bumble bees are selective about the floral resources they gather (Vaudo et al., 2015).

We find that crop pollination is maximised when two conditions are satisfied: First, the crop is located between the nest and the wildflower patch, and second, the floral resources offered by the wildflowers are significantly different from the resources offered by the crop flowers. This situation can occur, for example, if the crop flowers are deficient in certain nutrients that are present in the wildflower resources, and vice-versa. In addition, we find that only small quantities of wildflowers are needed to maximize crop pollination services when the composition of the wildflowers and the crop are similar, whereas large quantities are most beneficial when the wildflowers are very different in composition from crop flowers. We also showed how our predictions change as a function of the wildflower background density, which is a measure of the intensity with which the landscape is farmed. Intensively farmed monoculture landscapes are often nutritionally deficient (Girard et al., 2012; Topitzhofer et al., 2019). In our work, we find that the benefits of planting wildflowers adjacent to a crop increase as the agriculture becomes more intense. We emphasize that the benefits we found are not due to a pollinator population effect as our model does not include population dynamics, but only to the effect of the balance between nutrition and energy on pollinator spatial distributions. Häussler et al. (2017) pointed out that demographic effects can further augment the benefits of planting wildflower patches. Our work is an indication that spatial aspects of the system are also important. We therefore suggest that maximal effectiveness of wildflower patches can be achieved by combining spatial considerations with those aimed at supporting population dynamics.

## Acknowledgements

RCT acknowledge NSERC STPGP 506922-17 and NSERC DG RGPIN-2016-05277 grant. Also thanks BRAES and the BC Blueberry Council. EF acknowledge NSERC Discovery Grant.

